# The neural basis of emotional generalization in empathy

**DOI:** 10.1101/2025.10.18.683207

**Authors:** Jiayang Xiao, Anusha B. Allawala, Joshua A. Adkinson, Xiaoxu Fan, Melissa C. Franch, Victoria Gates, Raissa K. Mathura, Madaline Mocchi, John Myers, Bailey Pascuzzi, Suhruthaa Pulapaka, Garrett Banks, Eleonora Bartoli, Wayne K. Goodman, Sanjay J. Mathew, Xaq Pitkow, Nader Pouratian, Nicole R Provenza, Ben Shofty, Andrew J Watrous, Kelly Bjianki, Sameer A. Sheth, Seng Bum Michael Yoo, Benjamin Hayden

## Abstract

The essence of empathy is generalization of emotion across persons. Here, we leverage recent theoretical advances in the neuroscience of generalization to help us understand empathy. We measured brain activity in human neurosurgical patients performing two tasks, one focused on identifying their own emotional response and one identifying emotional responses in others. We quantified the representational geometry of local field potential (LFP) high-gamma activity in four regions: the medial temporal lobe, anterior cingulate cortex, orbitofrontal cortex, and insula. We found encoding of both self- and other-emotions in all four regions, but codes for emotion and person are disentangled (that is, factorized) in the insula, but not the other regions. This factorized representation allows for cross-person generalization of emotion in a way that tangled (non-factorized) representations do not. Together, these results support the hypothesis that the insula uniquely contributes to social mirroring processes by which we understand emotions across individuals.

## INTRODUCTION

The ability to empathize with other people requires *generalization* - that is, we must take structured knowledge gained in one context and apply it to a different one (Behrens et al., 2018; Tenenbaum et al., 2011). In the case of empathy, it requires understanding that the emotions we experience are similar to those experienced by other people (Happé et al., 2017; Preckel et al., 2018; Schurz et al., 2021). And, like other processes involving generalization, empathy requires us to simultaneously draw connections across contexts and maintain a separation between them. In the case of empathy, we need to understand that the emotion another person is feeling is similar to our own emotions while maintaining a self-other distinction, so that we don’t get confused and ascribe others’ emotions to ourselves. Failures to navigate this generalization process are a hallmark of many psychiatric diseases, including schizophrenia, depression, and autism (Fletcher-Watson & Bird, 2020; Gambin & Sharp, 2018; Nowacki et al., 2020). Consequently, the neural underpinnings of empathy have been of great interest in neuroscience and psychology (de Waal & Preston, 2017; Decety, 2015; Marsh, 2018).

Recent advances in neuroscience have led to the development of new theories about how the geometric coding properties of populations of neurons can promote generalization (Courellis et al., 2024; Nieh et al., 2021). Specifically, important work suggests that generalization relies on special representations in which population information is factorized, rather than conjunctive (Bernardi et al., 2020; Ostojic & Fusi, 2024). In conjunctive representations, information in different coding channels is entangled and thus not readily separable, while in factorized representations, the same information may be present but in a manner that is disentangled. In the context of empathy, brain systems that allow for cross-person transfer ought to have factorized representations of *emotion* and *person*.

The idea that empathy relies on shared neural representations that allow for abstracted, cross-personal emotional representation is supported by empirical data. For example, the ability to identify visceral states in others depends in part on circuitry related to perceiving our own emotions (Preckel et al., 2018; Singer & Lamm, 2009; Singer & Tusche, 2014). When people experience a painful stimulus, parts of the pain network, including the bilateral anterior insula and anterior cingulate cortex, are activated as when they see a loved one have a similar experience (Lamm et al., 2011; Singer et al., 2004). Likewise, multivoxel pattern analysis shows that feelings of disgust and perceptions of unfairness also co-activate the insula and cingulate cortex in similar ways (Corradi-Dell’Acqua et al., 2016). The overlapping activation is consistent with the hypothesis that observers infer the emotional states of others through a process that makes use of overlapping representations that help map the experiences of the observed onto the observers’ own state (de Waal & Preston, 2017). Indeed, some authors have conjectured that shared reactivations may be a core element of the theory of mind and may contribute to empathy as well (Mitchell & Phillips, 2015; Molenberghs et al., 2016; Schaafsma et al., 2015). However, while suggestive, we lack the critical evidence that the brain has factorized representations of self- and other-emotion.

Emotion activates much of the brain, including the medial temporal lobe (MTL), anterior cingulate cortex (ACC), orbitofrontal cortex (OFC), and insula (Hultman et al., 2016; Janak & Tye, 2015; Rajmohan & Mohandas, 2007; Underwood et al., 2021). Among these regions, the insula is of particular interest as a potential site for first-to-second-person transfer (Gasquoine, 2014; Menon & Uddin, 2010). The insula responds to viscerally relevant stimuli, such as pain or disgust-inducing odor, whether experienced personally or observed in others (Kanske et al., 2015; Lamm et al., 2011; Singer et al., 2009; Zhao et al., 2022). The insula is a key node in the salience network (Menon & Uddin, 2010; Uddin, 2017), which shapes our emotional experiences and contributes to the generation of emotional states (Cisler et al., 2019; Luo et al., 2014; Underwood et al., 2021). Moreover, insula contributes to the perception of bodily states, which in turn are thought to be foundational to emotion processing (Craig, 2002; Karnath et al., 2005; Paulus & Khalsa, 2021). Indeed, activity in the insula is elicited by a variety of emotional events, suggesting a role beyond visceral sensory processing (Nguyen et al., 2016; Uddin, 2015; Zhou et al., 2021). However, limited accessibility and the rarity of isolated lesions have provided difficult barriers to a more sophisticated understanding of its role (Molnar-Szakacs & Uddin, 2022; Uddin, 2017).

We recorded local field potentials in the human insula, as well as in MTL, ACC, and OFC. Participants performed two tasks, an emotional identification task (in which they identified their own emotional response) and an emotional expression identification task (in which they identified another person’s emotional response). In the insula, we found a response pattern in which emotions in both tasks were represented with a unique factorized geometry. Both forms of information were available in the other three regions, but the geometry was non-factorized, and thus inaccessible for transfer from first to second person. Thus, within our set of four sampled regions, cross-person generalizable representations appear to be unique to the insula. These findings suggest a possible site and mechanism for empathetic cognition. Moreover, they highlight the importance of coding properties for implementing functional differentiation between brain regions.

## RESULTS

### Task and recording sites

We recorded from intracranial electrodes while human epilepsy patients (n=17, nine female) viewed videos (*Emotional brain behavior quantification task*) and facial expressions (*Affective Bias task*) that varied along the emotional valence dimension (**Figure 1A**). The emotional brain behavior quantification (EBBQ) task was used to test for self-emotion, and the Affective Bias task was used for other-emotions. In the EBBQ task, stimuli consisted of 30-60 second duration emotional videos containing real-life news events (Samide et al., 2020). We used real-world news clips because they are clearly non-fictional, and may therefore provide greater validity than previous studies that used clips taken from movies or music videos (Abraham et al., 2008). Participants rated their own emotional valence at the end of each video. In the Affective Bias task, participants viewed a series of emotional facial expressions and rated the emotion of the face using a continuous scale by clicking on a slider bar (Bijanki et al., 2019; Metzger et al., 2023; Surguladze et al., 2004). The stimulus set consisted of computer-morphed faces spanning emotional valences and intensities (100% happy, 50% happy, 30% happy, 10% happy, neutral, 10% sad, 30% sad, 50% sad, and 100% sad).

**Figure 1:**
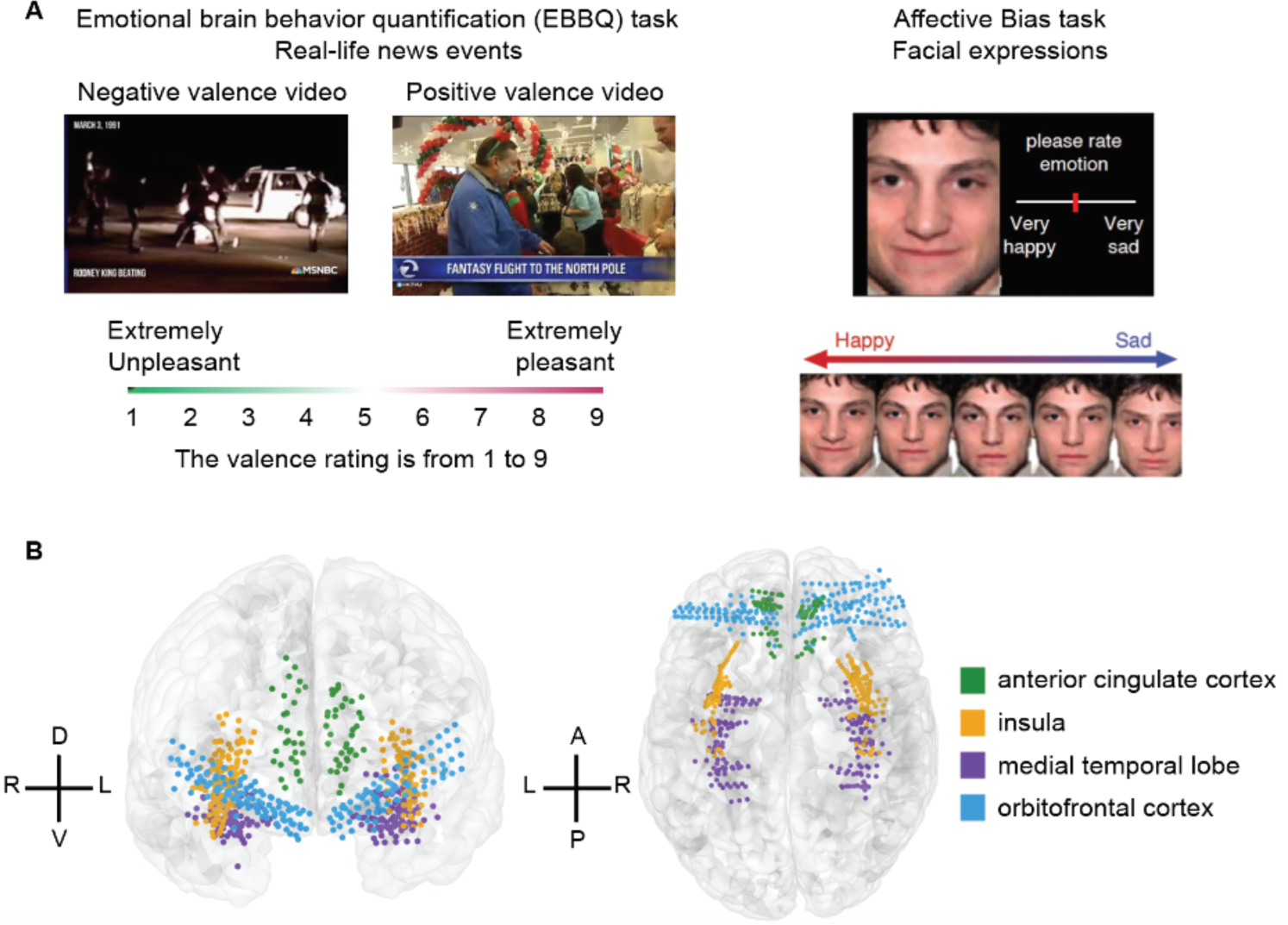
Self and other emotion tasks and neural recording locations. **(A)** Task structure. Each participant completed both EBBQ task (left) and Affective Bias task (right). The EBBQ task assessed self-emotion, where participants rated their own emotional valence at the end of each video using a keypad, ranging from 1 (extremely unpleasant) to 9 (extremely pleasant). The Affective Bias task evaluated other-emotion, where participants rated the emotion of a face by clicking a specific position on a slider bar. **(B)** Recording locations across all participants on a template brain. Coordinates are in Montreal Neurological Institute (MNI) space. Different colors indicate contacts from different regions: green, anterior cingulate cortex; orange, insula; purple, medial temporal lobe; blue, orbitofrontal cortex. Dorsal (D), ventral (V), left (L), right (R), anterior (A), and posterior (P) directions are indicated.

### Response to emotional valence is heterogeneous across contacts

We recorded 678 contacts from four brain regions in 17 participants who performed both tasks (**Figure 1B**). We first computed spectral power in gamma (35–50 Hz) and high-gamma (70–150 Hz) bands. Spectral power in these bands is a presumed index of local spiking activity from many neurons (Daume et al., 2024; Henrie & Shapley, 2005; Nir et al., 2007). Indeed, prior research has shown a close relationship between the spiking activity of neurons and the high-frequency activity observed in intracranial recordings (Lachaux et al., 2012; Nir et al., 2007; Rich & Wallis, 2017). Thus, it has long been proposed that spectral power in high-frequency bands, including gamma and high-gamma, may represent the collective activity of a local group of neurons and could potentially serve as a potential proxy for neuronal activity (Colgin et al., 2009; Fernández-Ruiz et al., 2021; Manning et al., 2009; Miller et al., 2014). We fit a linear regression model between spectral power and valence rating using the least-squares approach on the single contact level (**Figure 2A**). A positive normalized regression weight (shown in red) indicates the contact has a larger response in the gamma band during positive valence stimuli in each of the four regions we recorded (**Figure 2B-E**). These illustrative figures highlight the diversity of responses across contacts within each region.

**Figure 2.**
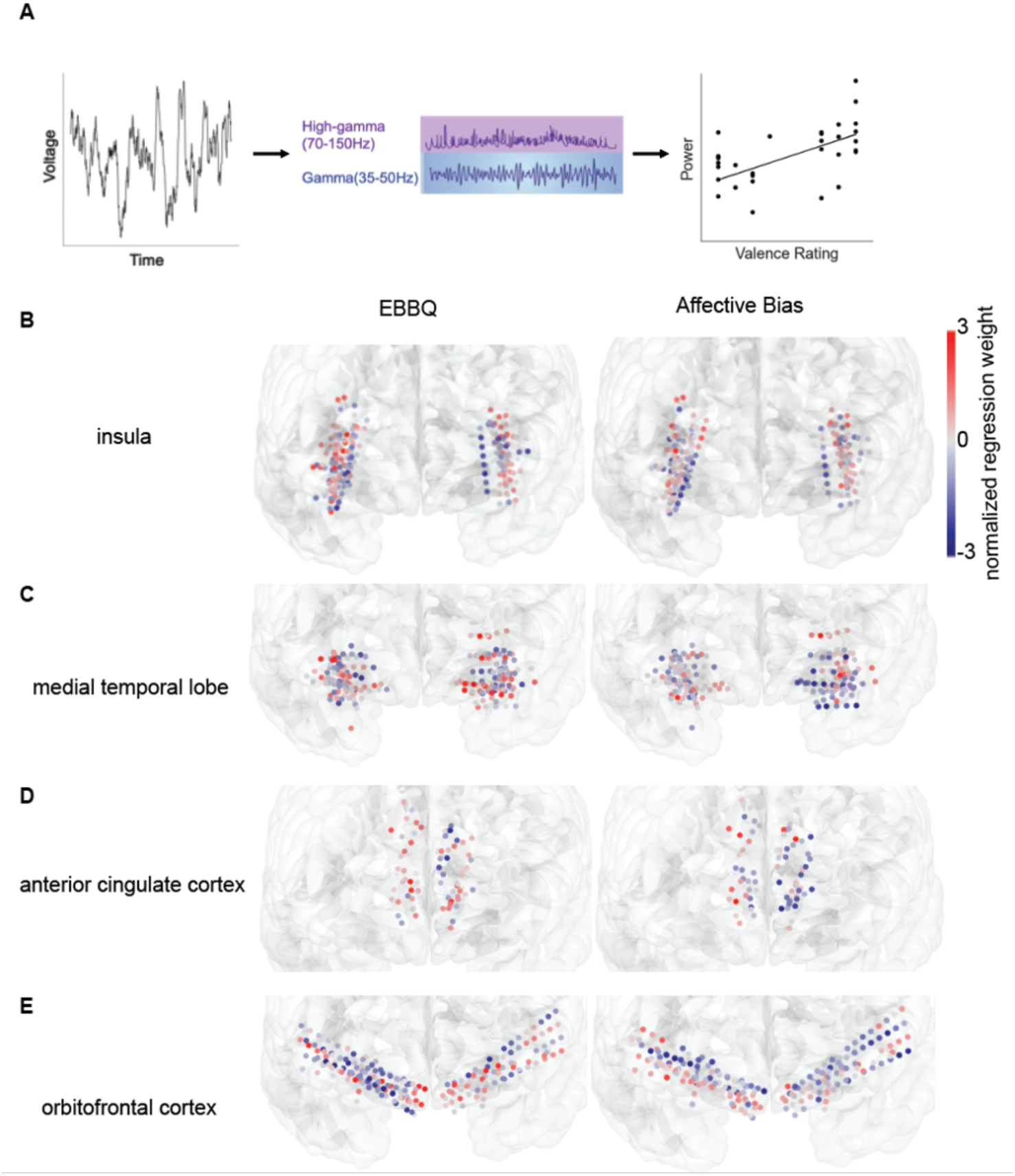
Neural response pattern across tasks. **(A)** For each contact, we obtained one normalized regression weight for each frequency band, representing the relationship between the spectral power in that band and the valence rating. **(B-E)** Normalized regression weight for gamma power across contacts during EBBQ and Affective Bias task. Contacts in red have a larger response during positive valence stimuli, while contacts in blue have a larger response during negative valence stimuli.

In the insula, we found clear encoding of emotional valence in both tasks. Specifically, the spectral power in gamma or high-gamma band was modulated by emotional valence in 13.5% of contacts during the EBBQ task, and 11.1% was modulated during the Affective Bias Task (EBBQ: n=28/208 contacts, Affective Bias Task: n=23/208 contacts, Bonferroni corrected because we checked two frequency bands). Both proportions were substantially greater than would be expected by chance (chance level: 5.0%, *p* < 0.05 in permutation test when compared to the true proportions). Valence was also encoded in both tasks in all three other regions we examined: the medial temporal lobe (MTL, EBBQ: n=29/198, 14.6%, Affective Bias Task: n=23/198, 11.6%), in the anterior cingulate cortex (ACC, EBBQ: n=11/69, 15.9%, Affective Bias Task: n=15/69, 21.7%), and in the orbitofrontal cortex (OFC, EBBQ: n=27/203, 13.3%, Affective Bias Task: n=29/203, 14.3%). These proportions from all regions were significantly larger than expected by chance (*p* < 0.05 in all cases). Overall, the proportion of contacts encoding emotional valence was similar between the EBBQ task and the Affective Bias Task. Indeed, we did not find any significant difference in the encoding of valence across tasks in all four regions (*p* > 0.9 for all four regions, chi-squared test). In both the EBBQ and the Affective Bias task, in all regions, the spectral power was greater during positive valence in some contacts, while in other contacts, the power was greater during negative valence; in other words, the response is heterogeneous across contacts (**Figure 2B-E**). Collectively, these results indicate that emotion is encoded in all four regions studied, consistent with findings from earlier studies (Jackson et al., 2024; Rogers-Carter et al., 2018; Rolls, 2019; Zheng et al., 2019).

### Generalized cross-person coding in insula but not other regions

To quantify the cross-person similarity of response patterns, we concatenated normalized regression weights from the two tasks and created vectors in which each entry corresponds to one recording site. The correlation between these two vectors tells us whether self- and other-emotion have similar effects on neural responses. In the insula, we found a positive correlation, indicating that the response to emotions is consistent across person (**Figure 3A**, gamma: r = 0.34, *p* = 4.24×10^-6^; high-gamma: r = 0.32, *p* = 2.79×10^-5^). We did not observe any even close to significant response in any other regions (**Figure 3B-D**, *p* > 0.9 for each region). Indeed, insula correlation coefficients were larger than those of other brain regions (Fisher’s z-transform, *p* values were Bonferroni corrected for gamma and high-gamma bands; compared with MTL: z = 4.19, *p* = 5.70×10^-5^ for gamma band, z = 2.59, *p* = 0.020 for high-gamma band; compared with ACC: z = 2.83, *p* = 0.0092 for gamma band, z = 3.04, *p* = 0.0047 for the high-gamma band; compared with OFC: z = 4.15, *p* = 6.77×10^-5^ for gamma band, z = 3.67, *p* = 4.81×10^-4^ for the high-gamma band).

**Figure 3.**
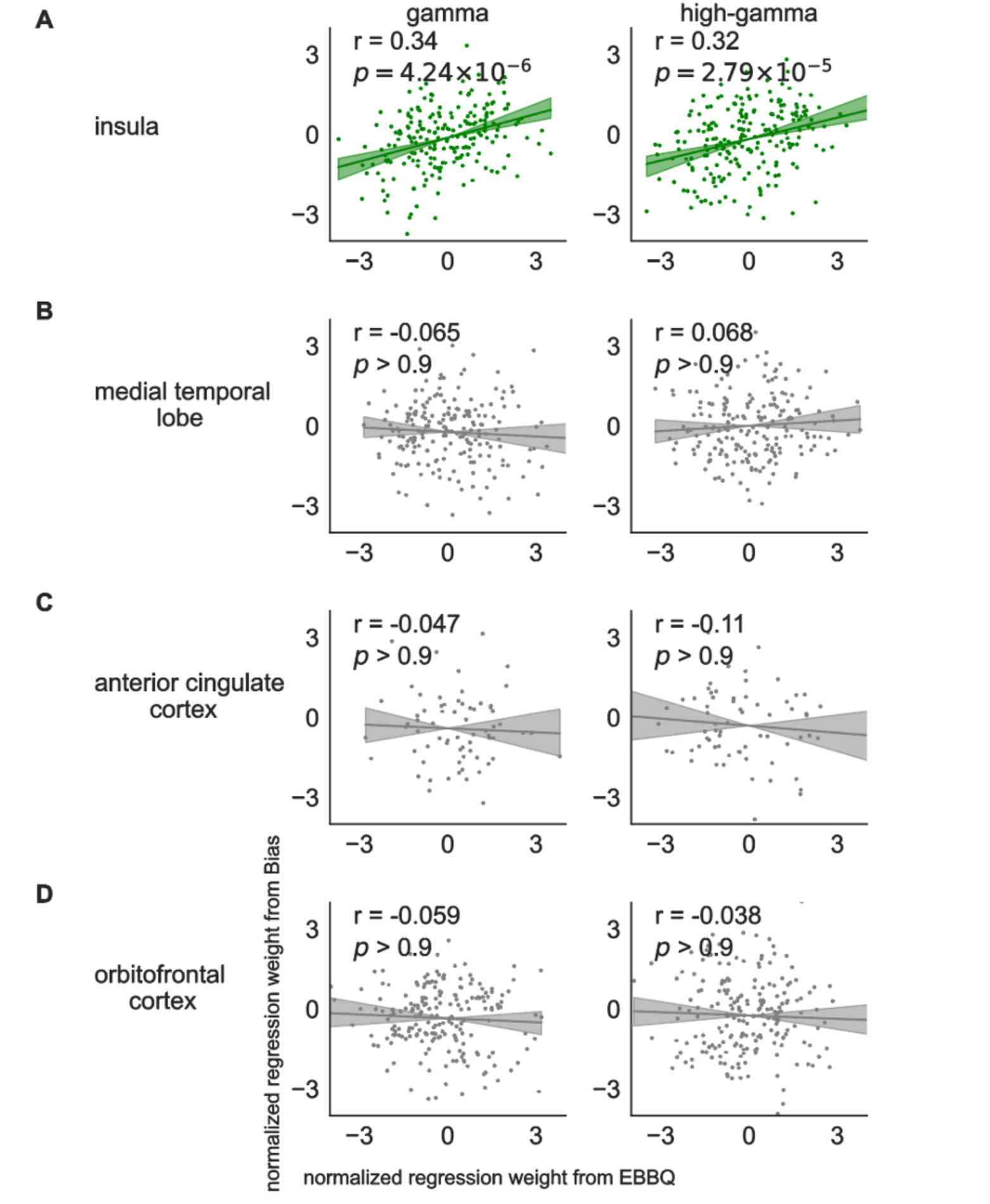
Cross-task coding in gamma and high-gamma activity. **(A)** Correlation between the normalized regression weights for gamma and high-gamma power from EBBQ task and the normalized regression weights from Affective Bias task. Each dot indicates one contact. **(C-D)** Correlation between the normalized regression weights from EBBQ task and Affective Bias task in other brain regions. Regions with significant positive correlations were shown in green while non-significant results were shown in gray.

We next assessed the extent to which the correlation between coefficients reflects abstract and task-specific codes using a positive-control / negative-control approach (Fine et al., 2023; Herman et al., 2023; Yoo & Hayden, 2020). We split the data within a single task into two halves and generated bootstrapped values from each half, providing an estimate of the correlation that would be measured given the noise properties of the stimuli under perfect correlation and repeated the entire process 1000 times (Azab & Hayden, 2017, 2018). In the medial temporal lobe, anterior cingulate cortex, and orbitofrontal cortex, the experimental values were significantly lower than the shuffled values (*p* < 0.05 in all cases). However, we did not observe a significant difference between these r-values in the insula (*p* > 0.5 in the insula). Thus, to the best of our ability to measure, insula coding of emotion does not contain a task-selective component and the other regions do not have a task-general component.

### Factorized representation of agency and emotion in human insula

We next asked whether *emotion* and *agency* are coded in a factorized manner in each of our four regions. We grouped data from positive-valence trials into one class and negative-valence trials into the other, ensuring an equal number of trials in each group. We combined trials from the self-task into one class and trials from the other-task into the other (**Figure 4A**). Using the spectral power in gamma and high-gamma bands, we decoded both valence (positive/ negative) and agency conditions (self/other). We found that the decoding accuracy of both agency and valence was higher than chance in the insula (**Figure 4B**, permutation test, *p(agency)* < 0.001, *p(valence)* < 0.001).

**Figure 4.**
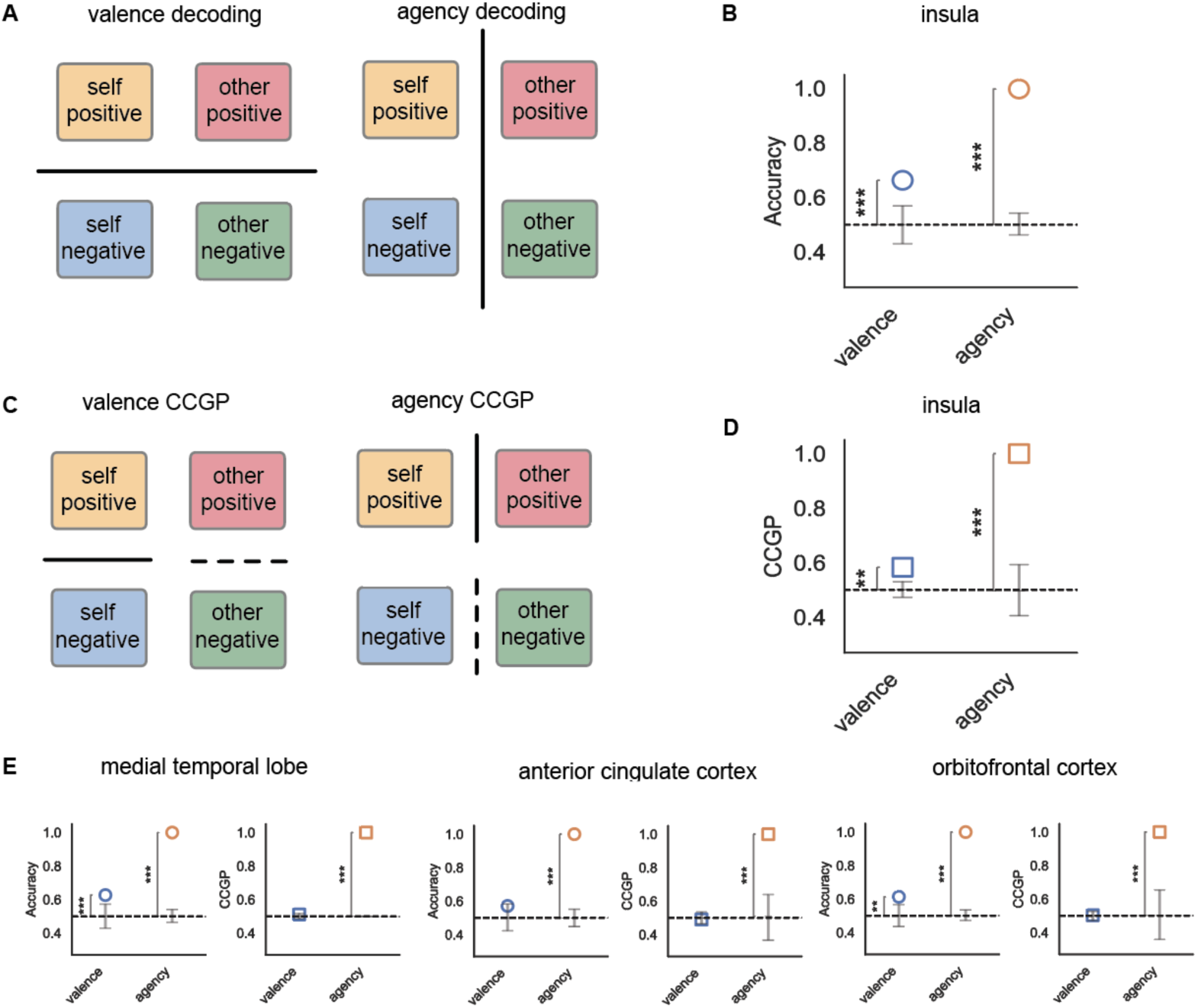
Factorized coding of emotion and agency. A) Decoding scheme for valence and agency. Decoding tests whether the information is present in a brain region. B) The decoding accuracy of valence and agency was significantly greater than chance in the insula. C) CCGP scheme for valence and agency. CCGP tests whether the information is represented in a disentangled manner and can generalize across conditions. D) CCGP of valence and agency was significantly greater than chance in the insula. E) Decoding accuracy and CCGP of valence and agency in the other regions.

We then assessed *coding geometry* using methods developed by Boyle et al. (2024). Specifically, we calculated the cross-condition generalization performance (CCGP) by training a classifier to decode valence using one task and testing using the other, then swapped the two sets and averaged the results (**Figure 4C**). We computed both valence and agency CCGP measures. We found that the CCGP of both agency and valence were higher than chance in the insula (**Figure 4D**, *p(agency)* < 0.001, *p(valence)* = 0.003). In other words, the decoder trained on one task can decode the valence in the other, consistent with a pattern of abstract valence encoding in the insula.

CCGP for the agency is significantly higher than the chance for all three regions, suggesting that the amount of information in each region is comparable (*p(agency)* < 0.001 for all regions). The decoding accuracy of valence is significantly higher than the chance for the medial temporal lobe and orbitofrontal cortex (medial temporal lobe: *p(valence)* < 0.001, anterior cingulate cortex: *p(valence)* = 0.07, orbitofrontal cortex: *p(valence)* = 0.002). However, the CCGP of valence is not significantly higher than chance in these regions (**Figure 4E**, medial temporal lobe: *p(valence)* = 0.07, anterior cingulate cortex: *p(valence)* = 0.79, orbitofrontal cortex: *p(valence)* = 0.25). Our results suggest that although valence is decodable in these regions, it is encoded in a factorized manner only in the insula.

### The coding axis of each emotion is correlated in the insula

We next examined whether the information axis representing each agent’s emotional valence was aligned or inversely oriented across the neural population. Even in the presence of parallel structure, an inverse orientation would imply that while decoding accuracy and CCGP scores could appear robust, a positive emotion in the self condition would correspond to a negative emotion in the other condition. To assess this, we first evaluated the high-dimensional geometric structure by calculating a parallelism score, which indicates alignment of the vectors encoding each variable. This score was significantly above zero in the insula (permutation test, *p* < 0.001 for valence, *p* < 0.001 for agency), but not in other regions (e.g., *p* > 0.5 for valence, **Figure 5A**). This suggests that the insula has a unique parallel structure representing the valence of emotion.

**Figure 5.**
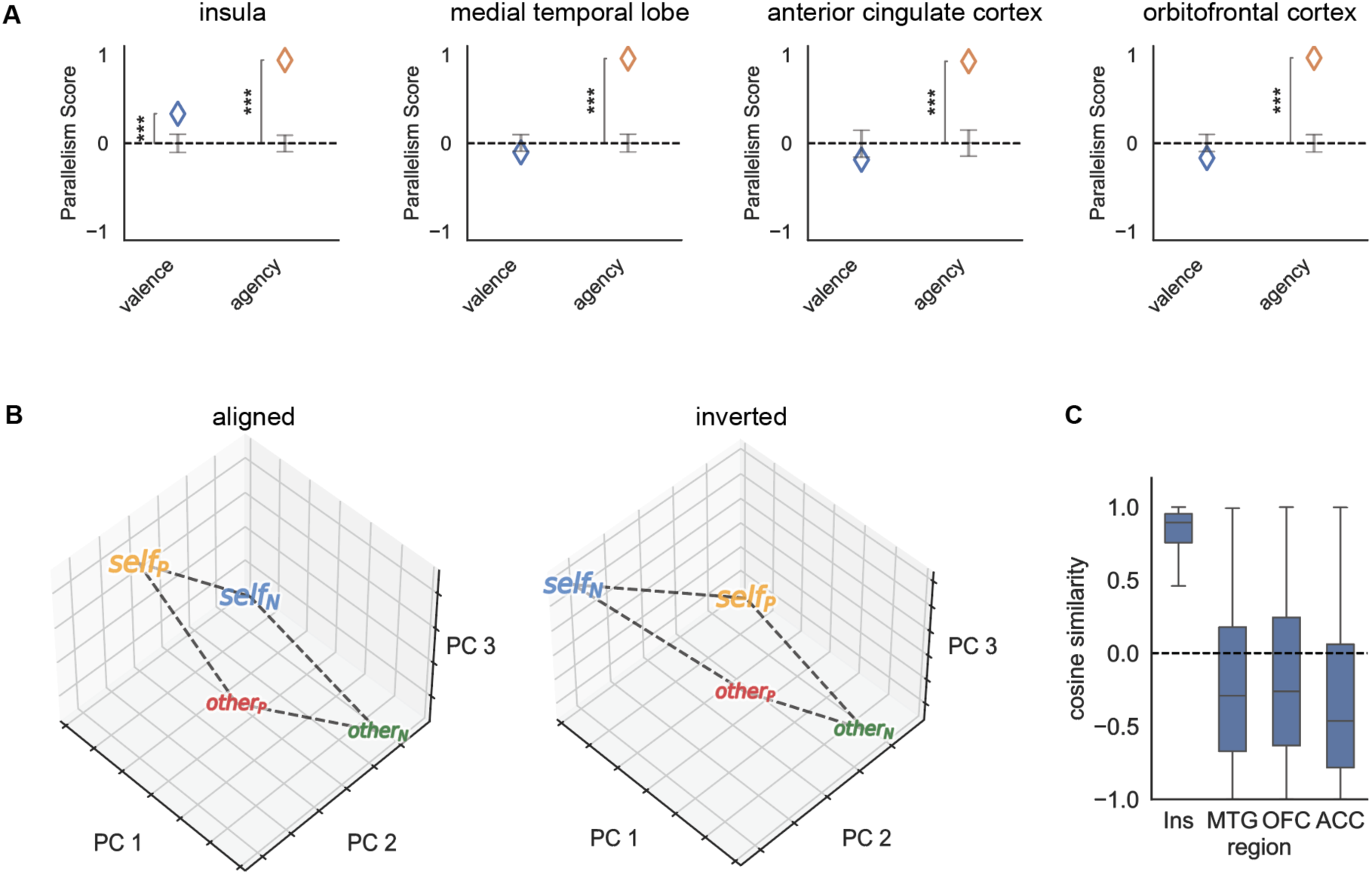
Alignment of the coding axis for emotional valence representation. A) Parallelism score across all four regions. A positive parallelism score indicates that the information axis representing the variable is aligned across conditions. B) Two possible cases for PCA projections of neural activity along the first three principal components. C) Cosine similarity between vector (self positive - self negative) and vector (other positive - other negative) across all regions.

We then applied principal component analysis (PCA) to the neural activity from four conditions—self-positive, self-negative, other-positive, other-negative. Projecting data into the space defined by the first three principal components could reveal whether the valence coding directions for self and other emotions were aligned or inversely oriented (**Figure 5B**). To quantify this alignment, we calculated the cosine similarity between the self-emotion vector (self-positive minus self-negative) and the other-emotion vector (other-positive minus other-negative). In the insula, the angle between these vectors was significantly aligned (angle = 35.9°, *p* = 0.013, **Figure 5C**), indicating that positive valence for the self condition aligns with positive valence in the other condition. In contrast, the vector angles for emotional valence were near orthogonal in other regions (medial temporal lobe: angle = 102.7°, anterior cingulate cortex: angle = 108.8°, orbitofrontal cortex: angle = 99.9°, *p* > 0.5 for all of these regions). These results suggest that the insula uniquely maintains an aligned coding axis for emotional valence across self and other contexts.

## DISCUSSION

We recorded intracranial field potentials while human neurosurgical patients performed a task in which they responded to emotional videos and, in a separate task, evaluated facial expressions for emotion. While information about self- and other-emotion was encoded in all four regions, the information was encoded in a ***factorized format*** in the insula but in a ***conjunctive format*** in the other three regions. The factorized coding format in the insula is the kind that would readily allow for generalizing emotional status from self to other (and vice versa, Bernardi et al., 2020; Ostojic & Fusi, 2024; Ruff et al., 2025). That is, it is a representational format that may allow us to draw conclusions about the emotions of others based on our own experience with those same emotions. We propose that these response patterns reflect an implementation of the computational processes that underlie empathy (de Waal & Preston, 2017; Preckel et al., 2018).

Historically, a dominant view held that arealization—the concept that specific brain regions are dedicated to specific functions—was the primary organizational principle of brain function (Posner et al., 1988). However, mounting evidence of distributed representations, often encapsulated in the phrase “everything is everywhere,” challenges this view, suggesting that variables of interest are broadly encoded across multiple structures (Steinmetz et al., 2019; Wang, 2022). This idea raises questions about how information is actually organized within the brain. One proposed mechanism for structuring information is *untangling*, where representations are organized to maximize linear separability and increase it hierarchically along the processing stream, supporting hierarchical encoding as information is distributed across regions (DiCarlo et al., 2012; Yoo & Hayden, 2018). Empirical and computational studies provide evidence for this untangling principle (Hénaff et al., 2019; Pagan et al., 2013; Ruff et al., 2025). Despite the widespread availability of information in latent form across many brain regions, our findings suggest that representational geometry differs between regions: some areas factorize information, while others maintain it in an entangled form. This observation offers a novel perspective on macroscopic computation, highlighting an objective beyond mere linear separability. Only discernible at the population level, this view reconciles arealization with distributed representation, suggesting that brain regions are not distinguished by the information they linearly access but by how they represent it. This differential representational geometry may be a fundamental principle in brain organization, moving beyond strictly localized or distributed models and providing insight into the specialized, region-specific roles of neural circuits.

A factorized or disentangled representation in neuroscience typically describes a representation scheme of the neural population where distinct factors in the abstract space correspond to separable, interpretable, and independent features of the stimulus or latent cognition (Ostojic & Fusi, 2024). Recently, factorized representation has been primarily identified in single-neuron resolution across species, notably in rodents and primates, and various cognitive domains (Bernardi et al., 2020; Boyle et al., 2024; Courellis et al., 2024; Johnston et al., 2024; Ruff et al., 2025; Okazawa et al., 2021; Chericoni et al., 2025). While this factorization approach provides insights into structured neural activity, our innovation demonstrates that similarly generalized, interpretable representations can be extracted directly from local field potentials (LFPs), in which each channel represents the aggregated activity of the neural population. As applying robust methods investigating population subspace and dynamics into macroscopic measurement offered insights for organization and computational principles (Kim et al., 2024), we hope the method of investigating factorized representation at the macroscopic brain activities would offer a scalable means to capture core principles of cognitive processes in humans.

These results are related to questions about how we recognize emotions in others and how the ability to do so relates to our emotional states (Ferretti & Papaleo, 2019; Spunt & Adolphs, 2019). Notably, in simulation-based theories, we can recognize emotional states in other people by stimulating the expression of emotions, that is, by recapitulating them (Goldie, 1999; Marsh, 2018). If this theory is correct, then it would help to have a part of the brain in which we have an abstract representation of the emotion so that it can be applied from self to other. Our results, therefore, suggest that the insula may be one such site in the brain.

The role of the insula in emotional processing remains a subject of ongoing debate (Gasquoine, 2014; Gogolla, 2017). The insula is activated by a wide range of emotional stimuli, from acute painful sensations to fearful facial expressions (Frot et al., 2022; Segerdahl et al., 2015). However, some researchers have proposed that the insula primarily serves as a sensory rather than affective region, with other regions, such as the orbitofrontal cortex, playing a larger role in value representation in subsequent processing stages (Rolls, 2016). A previous fMRI study, for instance, found that the anterior insula specifically encodes valence related to taste but not vision, whereas the orbitofrontal cortex encodes valence independent of the sensory source (Chikazoe et al., 2014). Our results challenge this idea.

To make our stimuli more relevant to real life, and thereby increase the ecological validity of our findings, we used videos containing non-fictional news clips as part of the study’s stimuli (Samide et al., 2020). However, using emotional videos and facial expressions may only partially capture the intricate nature of emotional processing. Emotion is a multifaceted phenomenon that involves various sensory modalities, including visual, auditory, tactile, and olfactory components (Barrett, 2006; Schirmer & Adolphs, 2017). Considering the involvement of the insula in gustatory and olfactory processing, as well as its role in multisensory integration, future research should consider incorporating additional sensory modalities to provide a more comprehensive understanding of how the insula processes emotions (Chikazoe et al., 2019; Gogolla, 2017; Koeppel et al., 2020).

The perception of emotional stimuli enriches our daily experiences and plays a vital role in our survival. Emotional valence processing could influence perceptual, attentional, and mnemonic processes, thus guiding our behaviors and shaping our interactions with the world around us (Ballhausen et al., 2015; Dolcos et al., 2020; Gutchess & Kensinger, 2018).

Disruptions in this crucial process have been strongly associated with various psychiatric conditions (Price & Duman, 2020; Tschida & Yerys, 2021). For example, neutral or ambiguous facial expressions tend to be evaluated as sad in individuals with depression (Bourke et al., 2010; Disner et al., 2011). This alternation in valence processing can profoundly impact how they perceive and interact with their environment. In the future, researchers can explore potential changes in neural activity among these patients. Understanding the neural underpinnings of emotional valence processing in healthy participants and individuals with psychiatric diseases may provide valuable insights for early detection and targeted treatment, ultimately offering important clinical benefits.

## METHODS

### Participants

Seventeen participants (eight males and nine females, mean age 43 years, range 19-63 years) undergoing invasive monitoring for the treatment of refractory epilepsy at Baylor St. Luke’s Medical Center (Houston, Texas, USA) participated in our study. The implantation sites were exclusively determined by the clinical team with the sole goal of localizing the seizure onset zone. Ethical approval for the study was granted by the Institutional Review Board at Baylor College of Medicine, under IRB protocol number H-18112. All participants provided both verbal and written consent to participate in the study. After the surgical implantation of electrodes, patients underwent approximately one week of inpatient monitoring. During this period, we conducted both the EBBQ task and the Affective Bias task (in the same patients) while concurrently recording neural activity. These tasks were presented on a Viewsonic VP150 monitor with a resolution of 1920 x 1080, positioned 57 cm away from the participants.

### Emotional brain behavior quantification (EBBQ) task

Each run in this task contained eight emotionally positive and eight emotionally negative videos. The valence category was predefined by average ratings from 100 healthy students in a publicly available database and all videos contained real-life news events (Samide et al., 2020). Unlike previous studies that have used video clips from movies or music videos, which might be processed differently due to their fictional nature, the video database employed in our research used real-life newscasts. At the end of each video, participants provided a valence rating using a USB numeric keypad, ranging from 1 (extremely unpleasant) to 9 (extremely pleasant). In this dataset, 16 out of the 17 participants completed two runs of this task, while the remaining participant completed four runs. Data from all runs were included in the analysis.

### Affective Bias task

Participants were asked to rate emotional human face photographs. We used a set of happy, sad, and neutral face examples (6 identities for each emotion, split evenly between genders), adapted from the NimStim Face Stimulus Set (Tottenham et al., 2009). These faces were manipulated through a Delaunay tessellation matrix to create subtle variations in emotional intensity, ranging from neutral to highly expressive. The increments were set at 10%, 30%, 50%, and 100% for happy and sad expressions, respectively.

In each trial, a white fixation cross was presented on a black background for 1000 msec (jittered +/- 100 msec). Then, a face image and a rating prompt were displayed simultaneously on the screen. The rating prompt included an interactive analog slider bar positioned under the text instruction: “Please rate the emotion”. Using a computer mouse, participants indicated their rating by clicking a specific position on the slider bar. The ratings were captured on a continuous scale ranging from 0 (’Very Sad’) to 0.5 (’Neutral’) to 1 (’Very Happy’). The presentation of stimuli followed a blocked design, with all happy faces (plus neutral) appearing in one block while all sad faces (plus neutral) appeared in a separate block. Each run consisted of one block of happy faces and one block of sad faces, with each block containing 30 randomized trials (6 identities x 5 levels of intensity). Participants completed between two to twelve runs with alternating happy and sad trials. All data from these runs were included in the analysis. The order of these blocks, either starting with happy faces or sad faces, was counterbalanced across participants.

### Electrode localization and intracranial recordings

Prior to surgery, patients underwent brain magnetic resonance imaging (MRI), followed by post-implantation computed tomography (CT). For precise localization of implanted electrodes, we aligned the pre-operative T1-weighted MRI scans with the post-operative CT scans. The automated cortical reconstruction was executed using Freesurfer tools (Fischl, 2012). After mapping the electrodes onto the native MRI space using BioImage Suite v3.5b1 (Joshi et al., 2011), an expert anatomist assessed the images to identify brain regions and categorized contacts as either gray or white matter. Contacts found within white matter were excluded from subsequent analyses. Detailed methodologies have been outlined elsewhere (Sheth et al., 2022; Xiao et al., 2023, 2024; Provenza et al., 2024 and 2025; Katlowitz et al., 2025). Neural signals were recorded using sEEG electrodes at a sampling rate of 2000 Hz through the Cerebus data acquisition system (BlackRock Microsystems, UT, USA). The signals were processed with a bandpass filter set at 0.3-500 Hz using a 4th-order Butterworth filter.

### Preprocessing and spectral power calculation

We conducted a visual inspection of the raw signals to identify any recording artifacts and interictal epileptic spikes. Contacts exhibiting excessive noise were excluded to prevent noise propagation to other contacts through re-referencing. To minimize the impact of line-noise artifacts and volume conduction, signals were notch-filtered (60 Hz and its harmonics) and then re-referenced through bipolar referencing (Bastos & Schoffelen, 2015). Subsequently, the referenced signals were down-sampled to 1000 Hz, followed by the application of a Hilbert transform to estimate spectral power across frequency bands. To determine spectral power values for each trial, we computed the average of squared magnitudes from the Hilbert transform decomposition. For the EBBQ task, spectral power was averaged across a time window beginning at video onset and ending at video offset. For the Affective Bias task, the mean spectral power was calculated across a time window beginning at the onset of the face stimulus and ending at the participant’s response.

### Data visualization in the template brain

Contacts were mapped onto the MNI space and visualized using the open-source software RAVE (R Analysis and Visualization of iEEG, Magnotti et al., 2020). To assess variations in neural responses to stimuli of different valence ratings, we used an ordinary least squares regression to generate one normalized regression weight for each contact. These normalized regression weights, which compared the average spectral power during positive valence stimuli with that during negative valence stimuli, were then plotted at each contact.

### Encoding of emotional valence in each task

A contact was considered selective for emotional valence when the spectral power in at least one frequency band was significantly modulated by valence in the linear regression model. To assess whether the proportion of selective contacts was significant, we derived a null distribution by repeating the same procedures after randomly permuting the valence ratings 1000 times. The p-value was defined as the probability that the proportion of selective contacts from the null distribution is higher than the true proportion. To test the encoding of emotional valence in both tasks, this analysis was conducted separately for each task in all four regions.

### Calculation of difference in correlation coefficient between regions

To compare the correlation coefficients, we transformed the correlation coefficients to Fisher’s z scores using the following equation for each correlation coefficient:

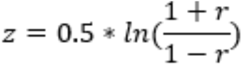

Then we computed the Z-test statistic for the difference between the two z scores using the following equation:

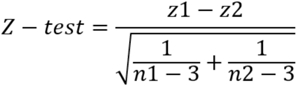

where z_1_ and z_2_ are the z scores transformed from the correlation coefficients for the two regions, and n_1_ and n_2_ are the sample sizes associated with the two values (number of contacts in each region).

Lastly, we determined whether the difference between the two correlation coefficients is statistically significant by comparing the Z-test statistic to the critical value from the standard normal distribution. A p-value of less than 0.05 indicates that the two correlation coefficients are significantly different from each other.

### Decoding performance

We conducted population decoding on a pseudo-population of channels pooled from all patients. Decoding was performed using a support vector machine with a linear kernel and default parameters in Decodanda (Boyle et al., 2024). To build the pseudo-population, we independently sampled trials for each channel, grouping them by the conditions for the decoding analysis. To decode valence, trials with positive valence were grouped as one class, while negative valence trials formed another. Similarly, to decode agency, self-task trials were categorized separately from other-task trials. The decoding accuracy is the mean accuracy across 1,000 iterations of random subsampling to mitigate variance from the sampling process.

### Cross-condition generalization performance (CCGP)

A key feature of neural representations of disentangled variables lies in their capacity to support generalization across conditions. To assess this, we used CCGP, which evaluates how well a decoder trained on one set of conditions can generalize to a separate set. To calculate CCGP, we held out trials from specific conditions during training, using only the remaining conditions to train the decoder, and then evaluated performance on the held-out conditions. For example, when decoding valence, we trained a classifier on trials from one task and tested on the other, then reversed the sets and averaged the results. When decoding agency, we trained a classifier to differentiate self-emotion from other-emotion using positive valence trials and tested on negative valence trials, then swapped and averaged as before. The final result is the mean CCGP across 1,000 iterations of random subsampling. To establish a null model for generalization, we needed to preserve the level of decodability while randomizing generalization. We used a rotation-translation of the pseudo-population vectors by shuffling channel indices and computed CCGP. We repeated this process to obtain a set of 1000 null CCGP values. The p-value for CCGP was defined as the probability that the CCGP from the null distribution is higher than the true CCGP.

### Parallelism score (PS)

To calculate the PS, we grouped trials into four conditions: self-positive, self-negative, other positive, other negative. The PS measures the correlation between coding directions, where each direction is defined by differences between conditions that vary only in the targeted variable. For example, to calculate the PS for valence, we computed the difference between self-positive and self-negative for one coding direction, and between other-positive and other-negative for the other direction. The PS was averaged over 1,000 iterations of random subsampling. We then shuffled channel indices to create a distribution of 1,000 null PS values. The p-value for PS was the probability that the value from the null distribution is higher than the observed value.

## Acknowledgments

We thank our patients for their participation in the research. This work was supported by the National Institutes of Health (UH3 NS103549 [to SAS, KRB, WKG, and NP], K01 MH116364 [to KRB], R01 MH127006 [to KRB], UH3 NS100549 [to WKG and SAS], and R01 MH114854 [to WKG]), McNair Foundation (to SAS, BYH, and NRP), and National Research Foundation of South Korea Grant No.IBS-R015-D1, RS-2023-00211018, and Ministry of Food and Drug Safety of South Korea Grant No. RS-2024-0033312 (to SBMY)

